# Forward-reverse mutation cycles between stages of cancer development

**DOI:** 10.1101/198309

**Authors:** Taobo Hu, Yogesh Kumar, Iram Shazia, Shen-Jia Duan, Yi Li, Lei Chen, Jin-Fei Chen, Rong Yin, Ava Kwong, Gilberto Ka-Kit Leung, Wai-Kin Mat, Zhenggang Wu, Xi Long, Cheuk-Hin Chan, Peggy Lee, Siu-Kin Ng, Timothy Y. C. Ho, Jianfeng Yang, Xiaofan Ding, Shui-Ying Tsang, Xuqing Zhou, Dan-Hua Zhang, International Cancer Genome Consortium, En-Xiang Zhou, Lin Xu, Wai-Sang Poon, Hong-Yang Wang, Hong Xue

**Author notes:** These authors jointly directed this work. Correspondence should be addressed to Hong Xue Division of Life Science Hong Kong University of Science and Technology Clear Water Bay, Hong Kong. These authors contributed equally to this work (Taobo Hu, Yogesh Kumar, Iram Shazia, Shen-Jia Duan).

## Abstract

Earlier, prominent occurrences of interstitial loss-of-heterozygosities (LOHs) were found in different cancers as a type of single-nucleotide-variations (SNVs), at rates far exceeding those of the commonly investigated gain-of-heterozygosities (GOHs) type of SNVs. Herein, such co-occurrences of LOHs and GOHs were confirmed in 102 cases of four cancer types analyzed with three different next-generation sequencing platforms, comparing non-tumor, paratumor, and tumor tissues with white-blood-cell controls; and in 246 pan-cancer cases of whole-genome tumor-control pairs. Unexpectedly, large numbers of SNVs enriched with CG>TG GOHs and copy-number-variations (CNVs) proximal to these GOHs were detected in the non-tumor tissues, which were extensively reversed in paratumors showing prominent TG>CG LOHs with proximal CNVs, and less so in tumors to form forward-reverse mutation cycles. Lineage effects in the reversions, likely resulting from directional selection, supported a sequential rather than parallel mode of evolution as described in a ‘Stage Specific Populations’ model of cancer development.

## Introduction

The progressive development of cancer has been investigated at the cytochemical and genetic levels^1-6^, and the existence of early premalignant stages in the process has been suggested by precancer manifestations in terms of DNA, gene expression, protein and microscopic changes^7-14^. Genomic analysis has played an increasingly important role toward delineation of mutational events^15,16^, and a recent study by us has led to the finding of massive single-base interstitial loss-of-heterozygosities (LOHs) in various types of cancers, pointing to widespread intersister chromosomal gene conversions arising from defective DNA double-strand break (DSB) repair in tumor development^17^. Thus the interplay between the gain-of-heterozygosity (GOH) and LOH types of single nucleotide variations (SNVs) requires examination along with copy number variations (CNVs) as mutational elements of cancer regarding the extent of possible reversion of forward mutations during cancer development.

In the present study, non-tumor, paratumor, primary tumor and metastatic tumor tissues in different types of solid tumors were compared with same-patient white-blood-cell controls based on massive parallel sequencing. Somatic mutations in both directions, *i.e.* GOHs, LOHs, CNV gains and CNV losses were examined residue-by-residue and window-by-window in order to detect the presence of mutation reversals in the development of cancer cells and to assess their biological significance. The results obtained suggested that forward-reverse cycles of mutations along with directional selection are important determinants of stage-specific cell population profiles in cancer development.

## Results

### Genotypic changes in non-tumor and paratumor tissues

White-blood-cells (B), tumor tissue (T), paratumor tissue (P) immediately adjacent to tumor, and more remote non-tumor tissue (N) were collected in twelve same-patient tetra-sample cases consisted of four breast carcinomas (BRCA), five stomach adenocarcinomas (STAD) and three hepatocellular carcinomas (LIHC) (Supplementary Table 1), and subjected to DNA analysis using the AluScan platform based on inter-Alu polymerase chain reaction (PCR) followed by massively parallel sequencing (MPS) as described in Methods. The genotype of a base residue was referred as a major allele (M) when it matched the genotype on human reference genome hg19, or as a minor allele (m) when there was no match, thereby enabling the identification of changes in the form of MM-to-Mm GOH (‘GOH-M’), mm-to-Mm GOH (‘GOH-m’), Mm-to-MM LOH (‘LOH-M’) or Mm-to-mm LOH (‘LOH-m’)^17^. Figs. 1a and 1b show the total number of residue-by-residue changes in the N-, P- or T-sample genomes relative to B-sample, *viz.* ΔNB, ΔPB or ΔTB respectively in terms of GOH-M, GOH-m and LOH (sum of instances of LOH-M and LOH-m). Since the numbers of GOH-M, GOH-m and LOH mutations were higher in ΔNB than in ΔTB, and comparable in ΔPB and ΔTB, both the N-sample and P-sample cells had to be regarded as premalignant or early malignant cells despite their normal morphology and expression of immunohistochemistry (IHC) markers, in contrast with T-sample cells showing enlarged nuclei (Fig. 1c) and reduced expression of IHC markers (Supplementary Table 1). Notably, a forward-reverse mutation cycle (FR-cycle) was commonly observed in the patch diagrams in Fig. 1b: GOHs were strongly reversed by LOHs (*e.g.*, G1 mutations reversed by L1 mutations), and LOHs strongly reversed by GOHs (*e.g.*, L14 reversed by G12). Moreover, the LOH mutations of Mm residues displayed pronounced lineage effects, favoring the formation of MM residues when the Mm residues themselves were derived from MM residues, but favored the formation of mm residues when the Mm residues were derived from mm residues (as in the LOH mutations highlighted by yellow triangles in the figure).

**Figure 1.**
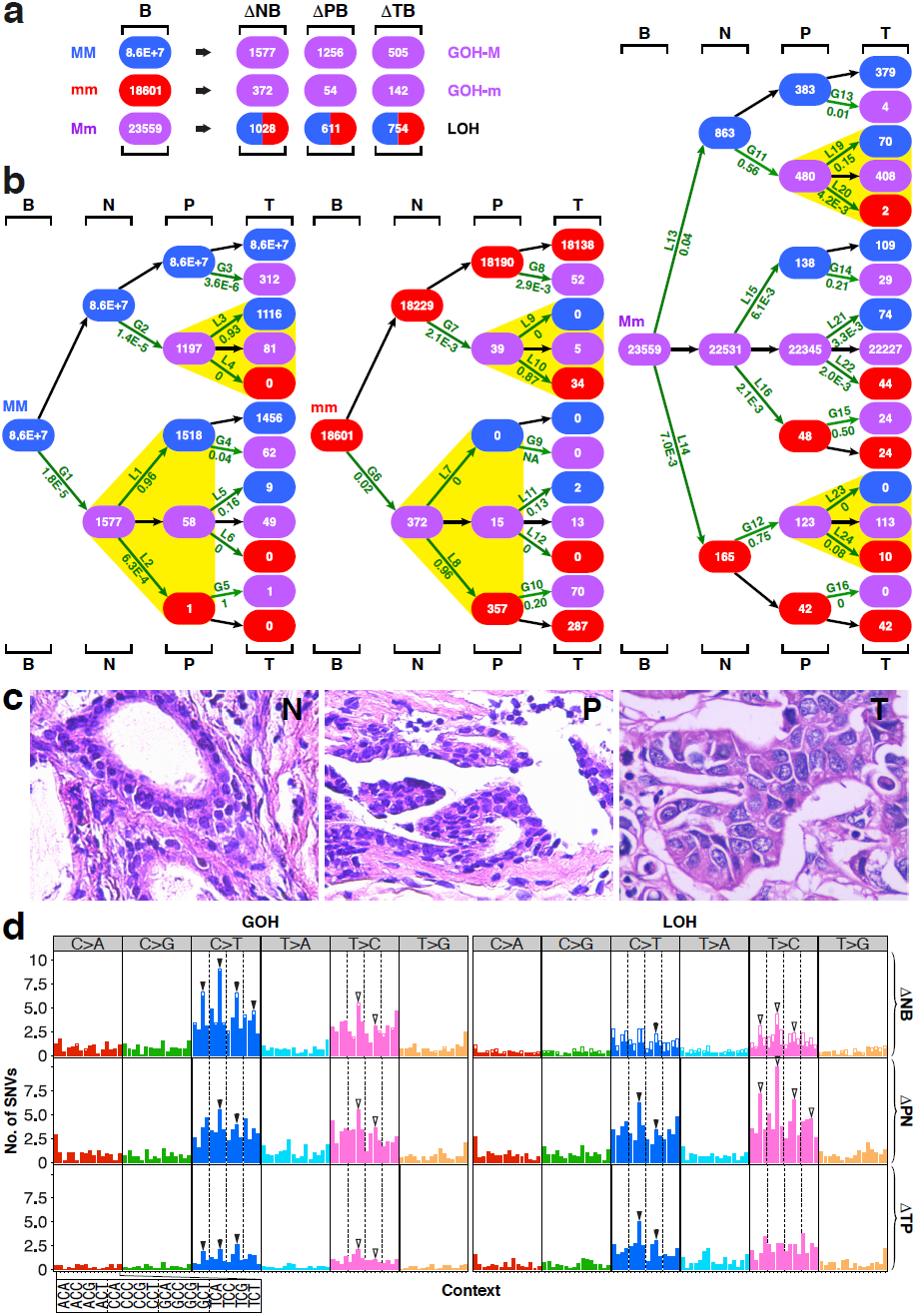
SNV mutations in B-N-P-T tetra samples. (**a**) Genotypic changes at N-, P- or T-stage of samples. The numbers of genotypic changes in N-, P- or T-stage sequences relative to B-stage sequences are represented by ΔNB, ΔPB and ΔTB respectively. LOH represents the sum of LOH-M and LOH-m changes. The twelve BN-P-T cases consisted of four breast carcinomas (BRCA), five stomach adenocarcinomas (STAD) and three hepatocellular carcinomas (LIHC) analyzed using AluScan sequencing (Supplementary Table 1). (**b**) Patch-diagrams tracing SNVs between the B-, N-, P- and T-samples originating from MM, mm or Mm residues in B-samples. Mutation rate is indicated below each LOH step (L1, L2, *etc.*) or GOH step (G1, G2, *etc.*). (**c**) Micrographs of N-stage tissue (left), P-stage tissue (middle) and T-stage tissue (right) in one of the representative BRCA B-N-P-T tetra samples. Magnification in each instance was 400. (**d**) Mutational profiles for the ΔNB, ΔPN and ΔTP SNV changes as numbered in the patch diagrams in Part **b**. The profiles are separated into the C>A, C>G, C>T, T>A, T>C and T>G types, where C>A includes both the C-to-A and the complementary G-to-T mutations, *etc*. Within each type, the 16 possible kinds of sequence contexts are indicated on an expanded scale on the x-axis, and the total number of SNVs observed for each kind of trinucleotide sequence contexts is represented by a vertical bar. In each vertical bar in the ΔNB tier, the solid segment represents the SNVs that were reversed in the next ΔPN tier, *e.g.* C>T GOHs being reversed by T>C LOHs, whereas the open segment represents the unreversed SNVs. Subgroups of contexts are compartmentalized by vertical dashed lines. Since residues of minor bases different from “m” were rare in the samples analyzed, mutations involving them are listed in Supplementary Tables 3 and 4 but not shown in the patch diagrams in Figs. 1b, 3b and 4b. M: major allele; m: minor allele; GOH-M:MM-to-Mm mutation; GOH-m: mm-to-Mm mutation; LOH-M: Mm-to-MM mutation; LOH-m: Mm-to-mm mutation.

Notably, in Fig. 1b right panel, partition of germline Mm residues via LOH steps L13 and L14 yielded a greater MM/mm product ratio than partition via LOH steps L15 and L16, and greater still than partition via LOH steps L21 and L22, although in each instance MM products exceeded mm products (Fig. 2a). Since all these three successive partitions emanated from germline Mm residues, their diminishing MM/mm ratios could not be the consequence of lineage effects. Instead, because MM residues in the genome generally have been optimized for growth by prolonged evolution, they tend to be favored over mm residues. Accordingly, the finding of [L13/L14] > [L15/L16] > [L21/L22] plausibly could be the result of a longer period of positive selection for MM over mm products of LOH in the N-stage cells relative to P-stage cells, and likewise P-stage cells relative to T-stage cells, suggesting that MM was generally preferred over mm for optimized growth of cells.

**Figure 2.**
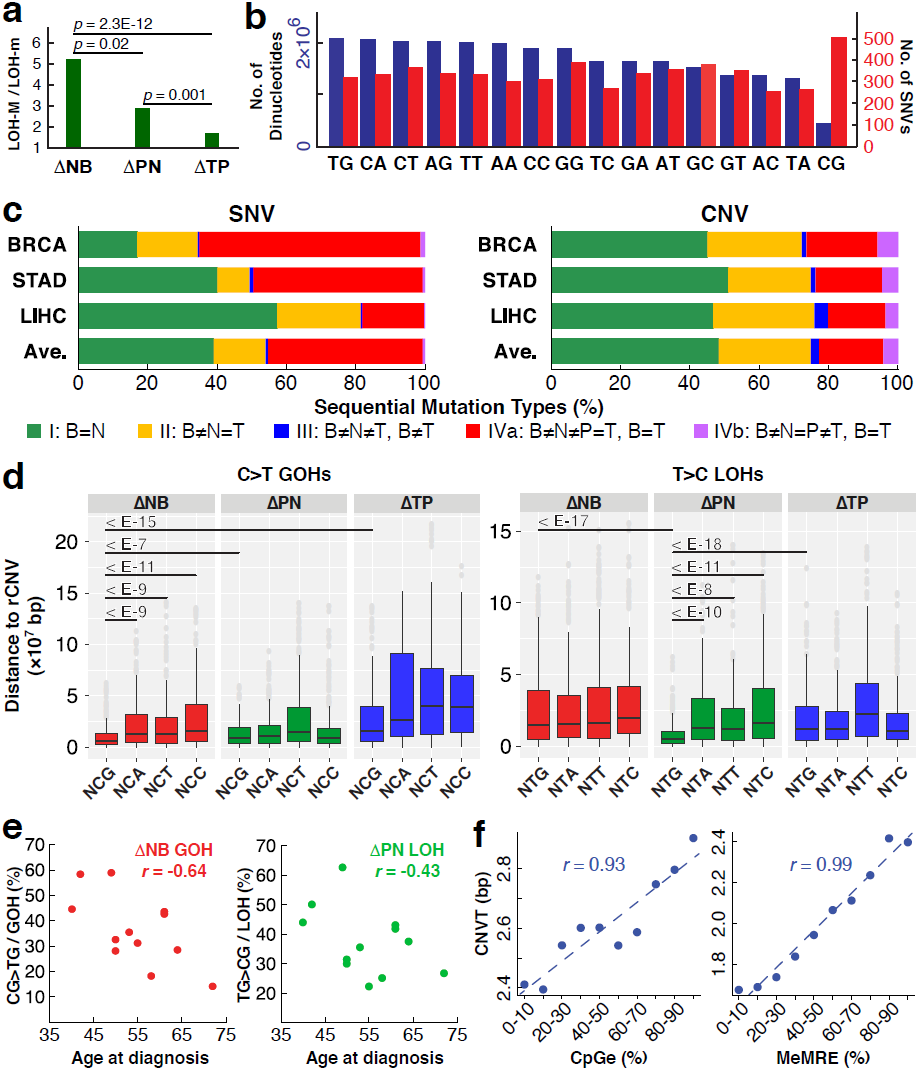
Properties of mutations in B-N-P-T tetra samples. (**a**) M-over-m preference in LOHs. LOH-M/LOH-m ratios for LOHs arising at germline Mm positions are shown for N-, P- and T-stages (data from Fig. 1b right panel). (**b**) Dinucleotides in genomic sequences captured by AluScan (blue bars), and SNVs found at the first base of dinucleotides (red bars). (**c**) Percentile of different types of sequential SNV and CNV changes in twelve tetra-sample cases of BRCA, STAD and LIHC. Type-I (green) B=N, *viz.* no SNV (or CNV) found in ΔNB. Type-II (yellow) B≠N=T, *viz.* same SNV (or CNV) found in ΔNB and ΔTB. Type-III (blue) B≠N≠T and B≠T, *viz.* altered in ΔNB and ΔTN, and also in ΔTB. Type-IVa (red) B≠N≠P=T and B=T, *viz.* altered in ΔNB and ΔPN, but not in ΔTP or ΔTB. Type-IVb (purple) B≠N=P≠T and B=T, *viz.* altered in ΔNB and ΔTP, but not in ΔPN or ΔTB. See Supplementary Tables 3 and 5 for detailed numbers of SNVs and CNVs at different stages. (**d**) Average distances between GOHs (left panel) or LOHs (right panel) and their nearest recurrent CNVs (rCNVs). Left panel shows that on average CG>TG GOHs among the ΔNB changes was closer to their nearest rCNVs than it was the case with eleven other kinds of C>T GOHs, with the five horizontal bars indicating how much closer in terms of *p* values. Likewise, right panel shows that on average TG>CG LOHs among the ΔPN changes was closer to their nearest rCNVs than it was the case with eleven other kinds of LOHs, with the five horizontal bars indicating how much closer in terms of *p* values. (**e**) Correlation between patient’s age at diagnosis and percentage of CG>TG GOHs among all GOHs for ΔNB (left panel), or percentage of TG>CG LOHs among all LOHs for ΔPN (right panel). (**f**) Correlation of numbers of somatic CNV breakpoints found in tumors (CNVT) from COSMIC database with either evolutionarily conserved CpG-rich regions (CpGe) (left panel), or unmethylated CpG-rich regions (MeMRE), from UCSC Table Browser database^22^ (right panel). *p*: significance probability; *r*: Pearson correlation coefficient (See Fig. 1 for abbreviations)

When the trinucleotide-based mutational profile method^18^ was employed to classify the GOHs and LOHs observed in the B-N-T-P tetra samples into the C>A, C>G, C>T, T>A, T>C and T>G groups, the results showed that C>T and T>C mutations were particularly prominent among both GOHs and LOHs, in keeping with the general expectation that transitions exceed transversions in SNVs (Fig. 1d). The C>T GOHs among the ΔNB changes displayed peak frequencies at the four NCG triplets, conforming to the ‘Signature 1A’ (marked by four solid arrowheads) common to cancers, and likely ascribable to the contribution of spontaneous deaminations of 5-methyl-cytosine at methylated CpG to form thymidine^18,19^. These deaminations would also explain the ˜50% greater occurrence of C>T GOHs than T>C GOHs in the ΔNB changes. In support of this, Fig. 2b shows that although there were less CG dimers than other dimers among AluScan captured as well as whole-genome sequences (Supplementary Fig. 1), more CG dimers underwent SNV mutations than any other dimers. In Fig. 1d, all SNV frequency columns in the ΔNB tier were represented by a solid segment and an open segment: the mutations in the solid segments were reversed in the next ΔPN tier, whereas the open segments were unreversed. Both the C>T and T>C GOHs show large solid segments indicating their extensive reversals in the ΔPN changes; since the T>C LOHs in the ΔPN tier were mostly reversal of the C>T GOHs in the ΔNB tier, these T>C LOHs were likewise more abundant than ΔPN C>T LOHs, and showed four NTG peaks (marked by open-arrows), which may be referred to as a ‘Signature 1A’ LOH feature.

Fig. 2c summarizes the forward and reverse mutation occurrences in the B-N-P-T samples: more SNVs and CNVs occurred in N-stage (*viz.* sum of Types II, III, IVa and IVb patterns) than in P- and T-stages combined (*viz.* Type-I). Reversals of N-stage SNVs and CNVs (*viz.* sum of Types IVa and IVb patterns) were common, amounting to ˜70% of N-stage SNVs or ˜40% of N-stage CNVs; and far more of such reversals took place in P-stage (Type-IVa) than in T-stage (Type-IVb).

When another seventeen B-N-T trio sample sets consisted of one BRCA, two LIHC and fourteen non-small cell lung cancers (NSCLC) were analyzed with respect to GOH and LOH changes in the N- and T-stage cells relative to B-stage cells (Fig. 3), the results obtained showed the same regularities as the B-N-P-T tetra samples: the B-genomes displayed much higher LOH (L5, L6) rates and GOH-m (G3, G4) rates than GOH-M (G1, G2) rates, strong lineage effects in LOH-partitions between MM and mm products (highlighted by yellow triangles), and prominent FR-cycles *viz.* L1 reversing G1, L4 reversing G3, G5 reversing L5, and G6 reversing L6.

**Figure 3.**
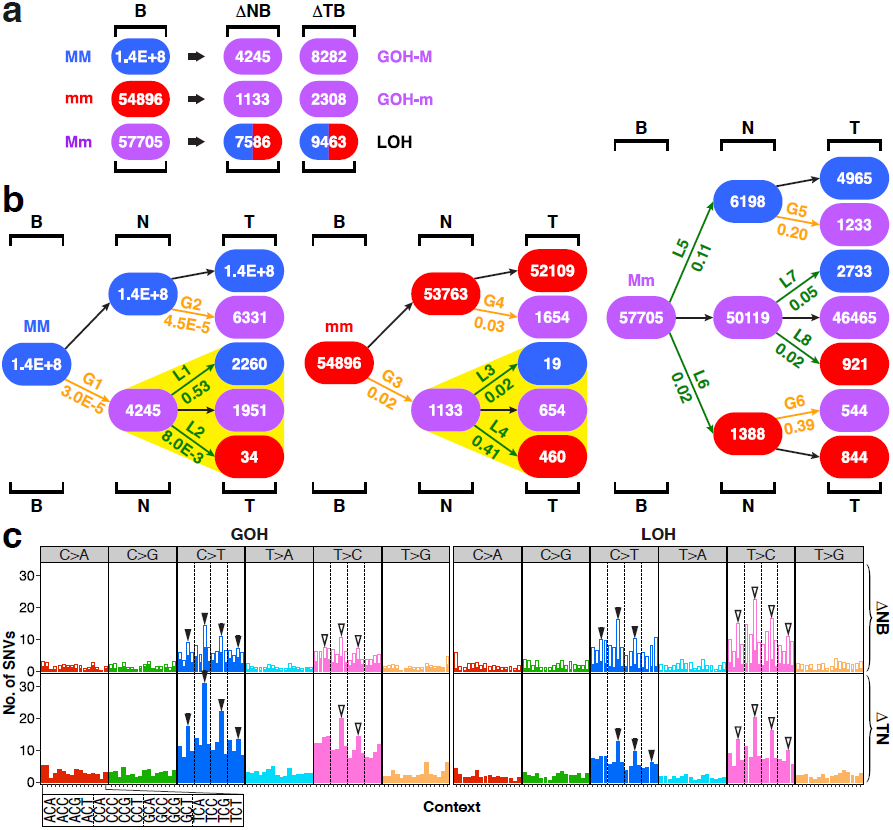
SNV mutations in B-N-T trio samples. (**a**) Genotypic changes in N- or T-stage of samples. The numbers of genotypic changes in N- or T-stage sequences relative to B-stage sequences are represented by ΔNB and ΔTB respectively. The seventeen B-N-T trio samples consisted of one BRCA, two LIHC and fourteen non-small cell lung cancer (NSCLC) cases. (**b**) Patch-diagrams tracing SNVs between the B-, N- and T-samples. (**c**) Mutational profiles for the ΔNB and ΔTN SNV changes as numbered in the patch diagrams in Part **b**. In each vertical bar in the ΔNB tier, the solid segment represents the SNVs that were reversed in the ΔTN tier, whereas the open segment indicates the unreversed SNVs. (See Supplementary Tables 6 and 7 for details of SNVs, and Fig. 1 for abbreviations)

### Genotypic changes in primary and metastatic tumors

Figs. 4 and 5 compare the mutations observed in five cancer groups based on same-patient N-stage, T-stage and metastatic-stage (M-stage) samples: (i) AluScan group of two N-T-M trio sets analyzed with AluScan sequencing; (ii) WGS-Liver-M group of four trio sets of liver-to-lung metastasis analyzed by Ouyang *et al*^20^ using whole genome sequencing (WGS); and sixty-seven trio sets involving brain metastases analyzed with whole exome sequencing (WES) by Brastianos *et al*^21^, which were separated into (iii) thirty-eight WES-Non-Lung cancers, (iv) six WES-NSCLC-L (L = low in C>A GOHs) cancers and (v) twenty-three WES-NSCLC-H (H = high in C>A GOHs) cancers. Although the five N-T-M trio groups compared in Fig. 4a were analyzed using variously the AluScan, WES and WGS platforms, the ratios of the [ΔTN]/N and [ΔMN]/N counts both indicated that the rates of LOH far surpassed the rates of GOH-m, which in turn far surpassed the rates of GOH-M. All five groups also displayed pronounced lineage effects in Fig. 4b in the partitions of LOH mutations of Mm residues between MM and mm products (highlighted by yellow triangles).

In Fig. 5a, the relative prominences of ΔTN GOHs, ΔTN LOHs, ΔMT GOHs and ΔMT LOHs varied among the five different cancer groups. This could arise in part from biological dissimilarities between the sequences analyzed on the different platforms on account of their varied sequence coverages of the genome. The SNV sites observed in the five groups displayed non-identical distributions among the Genic, Proximal and Distal sequence zones^22^, as well as non-identical replication timings during the cell cycle (Fig. 5b). The proportion of ΔTN GOHs that became reversed in the ΔMT changes, marked by solid segments of the GOH frequency bars in the ΔTN tiers, were highest in the AluScan group, also quite high in the WESNSCLC-L group, modest in the WES-Non-Lung group and lowest in the WGS-Liver-M and WES-NSCLC-H groups, even though the WES-NSCLC-L, WES-Non-Lung and WES-NSCLC-H groups were all analyzed based on the WES platform^21^.

**Figure 4.**
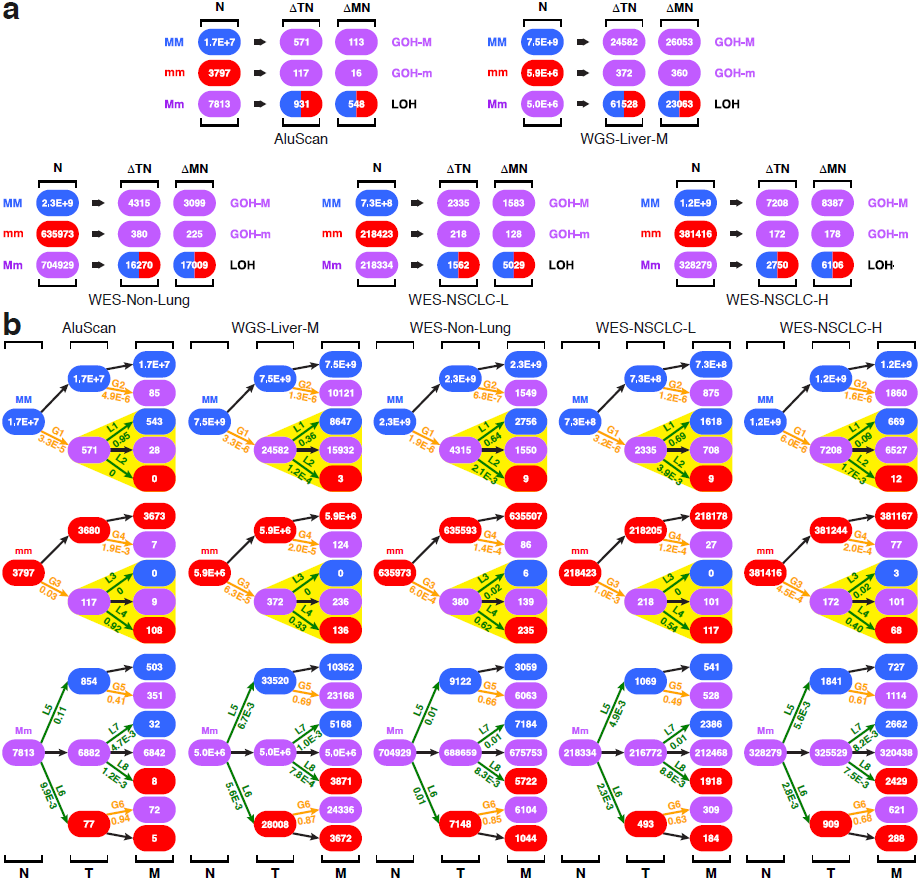
SNV mutations in N-T-M trio samples. (**a**) Genotypic changes in T- or M-stage sequences. The numbers of genotypic changes in T- or M-stage sequences relative to N-stage sequences in each of five case groups are represented by ΔTN and ΔMN respectively. The five case groups include AluScan group of two N-T-M trio sets analyzed with AluScan sequencing; WGS-Liver-M group of four trio sets of liver-to-lung metastasis analyzed by whole genome sequencing (WGS); and sixty-seven trio sets involving brain metastases analyzed with whole exome sequencing (WES) which were separated into thirty-eight WES-Non-Lung cancers, six WES-NSCLC-L (L = low in C>A GOHs) cancers and twenty-three WES-NSCLC-H (H = high in C>A GOHs) cancers. In the trio sets of WGS-Liver-M, WES-Non-Lung, WES-NSCLC-L and WES-NSCLC-H, because non-tumor tissues were sampled at > 2cm from tumor’s edge as controls instead of blood cells, the samples were designated as ‘N-stage’ for comparability with Figs. 1a and 3a. (**b**) Patch-diagrams tracing SNVs between the N-, T- and M-samples. (See Supplementary Tables 8 and 9 for details of SNVs). M: metastatic tumor. (Liver = hepatocellular carcinoma; NSCLC = non-small-cell lung cancer; Non-Lung = all 12 types of solid tumors in Ref. 21 except lung adenocarcinomas and lung squamous carcinomas; See Fig. 1 for abbreviations)

**Figure 5.**
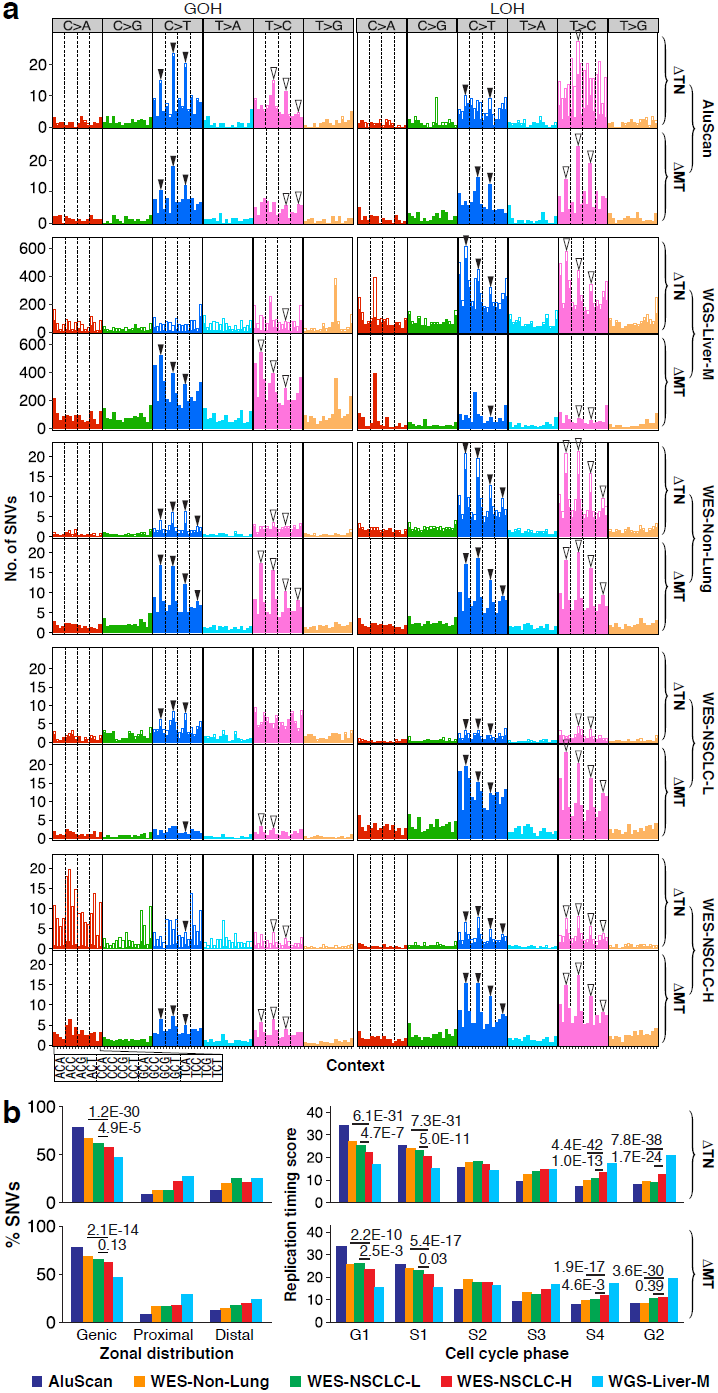
Properties of SNVs in N-T-M trio samples. (**a**) Mutational profiles for the ΔTN and ΔMT SNV changes as numbered in the patch diagrams in Fig. 4b. In each vertical bar in the ΔTN tier, the solid segment represents the SNVs that were reversed in the ΔMT tier, whereas the open segment indicates the unreversed SNVs. (**b**) Zonal distribution and replication timing of the SNVs in the ΔTN tier and the ΔMT tier in five case groups. Left panels: zonal classification of SNV (GOH and LOH) sites determined as described^22^. Right panels: replication timing scores derived from ENCODE at UCSC based on the “Repli-chip” method^47^. (See Figs. 1 and 4 for abbreviations)

The WES-NSCLC-H group was unique in its display of particularly eminent C>A GOHs. Previously, C>A transversions were linked to polycyclic aromatic hydrocarbons^23^ and acrolein^24^ in tobacco smoke. The 23 WES-NSCLC-H samples were derived entirely from smokers, in accord with smoking being a significant factor for their elevated C>A GOHs. However, the WES-NSCLC-L samples with much more subdued C>A GOHs included two non-smokers and four smokers, suggesting that smoking or high C>A GOHs could play a less important carcinogenic role in a minority of smokers.

### Whole genome sequencing confirmed the abundance of interstitial LOHs

A total of 246 tumor-control pairs from International Cancer Genome Consortium (ICGC) collection of WGS data^25^ were analyzed to yield LOH and GOH types of SNVs in each paired-samples. These included a panel of sixty-three pan-cancer cases (Pilot-63) (Fig. 6a), and twenty-two intrahepatic cholangiocarcinoma (ICC), eighty-six LIHC and seventy-five NSCLC cases (Fig. 6b), showing prominent LOHs in each instance. For the Pilot-63 dataset, the different ΔTB mutation counts in T-stage cells relative to B-stage cells (Fig. 6a left panel) yielded a rate ratio of 4,300 between LOH and GOH-M, which was comparable to the 5,400, 2,700 and 5,300 rate ratios observed in Fig. 1, Fig. 3 and earlier in Ref. 17 respectively, indicating a vastly greater rate of LOH than GOH in the cancer cells in all four instances. As well, in all four instances, the three mutation rates remain in the same order of R_LOH_ > R_GOH-m_ > RGOH-M, with LOH rate being the highest. The massive interstitial LOH rates observed earlier based on AluScan data were thus confirmed by the ICGC Pilot-63 WGS dataset.

**Figure 6.**
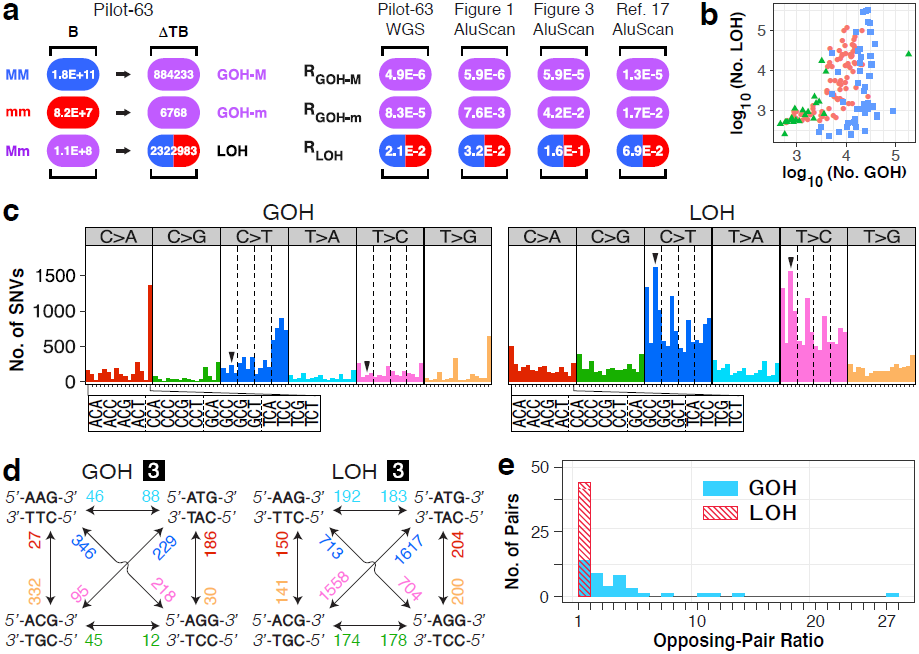
Properties of ICGC-whole genome sequencing of tumor (T)-blood (B) pairs. (**a**) Left panel: numbers of different genotypic changes between T- and B-stage sequences (ΔTB) in ICGC Pilot-sixty-three pan-cancer samples. Right panel: rates of different types of SNV mutations. For Pilot-63 samples, the rates were derived from WGS ΔTB/B values in the left panel; for B-N-P-T AluScan samples, from ΔTB/B values in Fig. 1a; for B-N-T AluScan samples, from ΔTB/B values in Fig. 3a; for Ref. 17 samples, from AluScan results on six types of cancers^17^. (**b**) Scatter plot of numbers of GOHs and LOHs in twenty-two intrahepatic cholangiocarcinoma (green triangles), eighty-six LIHC (red circles) and seventy-five NSCLC (blue squares) pairs of samples (Supplementary Table 10). (**c**) Alteration-group plots of mutational profiles for GOHs and LOHs between B- and T-stages in Pilot-63 samples. Each plot was grouped using the six alteration types: C>A, C>G, C>T, T>A, T>C, and T>G. (**d**) Representative mutation-rate diagrams, GOH-3 (left panel) and LOH-3 (right panel), were generated from the ICGC Pilot-63 dataset (See additional nine pairs of diagrams in Supplementary Fig. 2). In these diagrams based on triplet duplexes, the rates of opposing mutations from Part **c** were paired and labeled on bidirectional arrowheads with same color-codes as in Part **c**, *e.g.*, the rates of ACG>ATG GOH and ATG>ACG GOH changes (arrowhead-marked in Part **c** left panel) were dissimilar (229 in blue versus 95 in pink), but the rates of LOH changes of the same triplet duplexes were similar (1617 in blue versus 1558 in pink, and arrowhead-marked in Part **c** right panel). (**e**) Distribution of the observed rate ratios between opposing GOH pairs (the blue columns) and LOH pairs (the striped red column). (See Fig. 1 for abbreviations)

### Evidence from mutational profiles for gene conversions in LOH production

For the ΔTB SNVs of Pilot-63 (Fig. 6a), the alteration-group plots of mutational profile in Fig. 6c revealed strikingly similar heights for the opposing C>T (blue) and T>C (pink) LOH bars in the right panel, but not for the opposing C>T (blue) and T>C (pink) GOH bars in the left panel, consistent with excess LOHs than GOHs. The context-group plots in Supplementary Fig. 2a show comparable rates for all pairs of opposing LOHs in contrast to the unequal rates for the opposing GOHs. The mutation-rate diagrams in Fig. 6d show the actual rates for all opposing GOH pairs and LOH pairs. These results are summarized in Fig. 6e, where over 61% of the rate ratios of opposing substitutions were greater than two for the GOHs (blue bars), but 100% were between one and two for the LOHs (striped red bars), supporting different underlining mechanisms for GOH and LOH. This divergence of the rate ratios between opposing GOHs and opposing LOHs was in accord with our proposal that the LOHs in cancer cells resulted mainly from DSB repairs through gene conversions^17^. In a DSB at a heterozygous C/T, gene conversion could yield either a C/C or T/T homozygous position at comparable rates, depending on which homologous chromatid bears the DSB. On the other hand, because GOHs depend on point mutations rather than gene conversions, this comparable-rate constraint would not apply to GOHs.

### Distances between SNVs and recurrent CNVs

That the N-stage SNVs and CNVs in the B-N-P-T tetra samples both underwent active reversions, and more in P-stage than in T-stage (Fig. 2c) suggest some form of possible correlation between these two types of mutations. This was supported by Fig. 2d which shows that the sites of C>T GOHs with NCG context occurring in the ΔNB changes, and T>C LOHs with NTG context occurring in the ΔPN changes, were located particularly close to recurrent CNVs compared to mutations with other contexts or in other stages of change, *p* < 10^-7^. Furthermore, these two groups of SNVs declined in old age (Fig. 2e), in resemblance to the decrease of global DNA methylation in old age^26^. The correlation between somatic CNVs with CpGe and MeMRE (Fig. 2f), the increased SNVs at CpG sites (Fig. 2b) and the high tendency of methylated CpG conversion to TpG^27^, also pointed to some SNV-CNV relationships in the CNV process merits investigation.

### Frequency classes of serial CNV changes

In the B-N-P-T tetra-sample cases, the status of any CNV in the N-, P- and T-stages could be CN-unaltered (U), CN-gain (G) or CN-loss (L) relative to its status in B-stage. Arranging in serial order, the CN-status found in the N-P-T stages (Supplementary Tables 5) yielded twenty-six different serial orders, and their frequencies fell into three classes (Figs. 7a and 7b). In the LUG order, for example, each CNV site was CN-loss in N-stage, CN-unaltered in P-stage and CN-gain in T-stage, and the total number of sequence windows in the B-N-P-T sequences analyzed that exhibited such an LUG order made up the frequency on the y-axis of Fig. 7a. The three frequency classes separated by vertical dashed lines in the Fig. were:

I. *Class I (U=2)*: comprising six different orders, where a U status occurred in two of the N-, P- and T-stages.
II. *Class II (U≤1)*: comprising eight different orders, where the U status occurred in no more than one of the N-, P- and T-stages.
III. *Class III (Disadvantaged)*: comprising twelve different orders, where ten of them (*viz.* outside of LUG and GUL) included an abrupt double-dose change directly from G to L, or L to G in the order.

**Figure 7.**
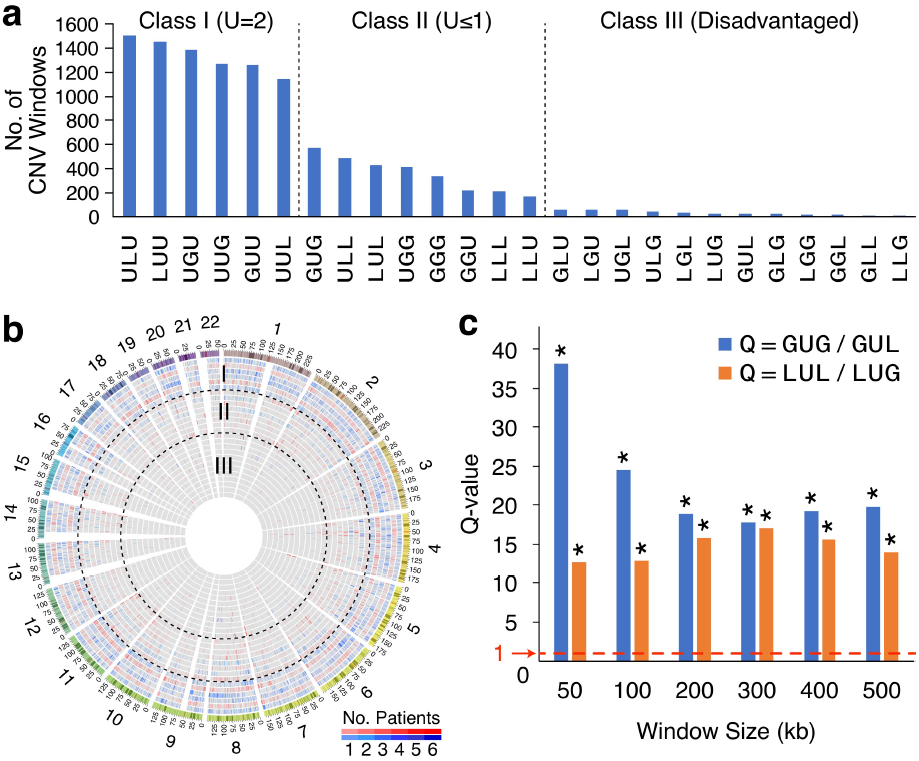
Serial orders of copy number (CN) changes in B-N-P-T tetra samples. (**a**) Frequencies of three classes of sequential orders of CN changes. Total numbers of CNV windows conformed to each serial order of changes are plotted out for all twenty-six possible orders. The CN-status of 500-kb windows of N-, P- or T-stage was determined relative to B-stage for twelve B-N-P-T tetra samples as described in Methods. The total numbers of windows that conformed to the different possible serial orders of CN changes (Supplementary Table 11) provided the basis for their partition into three frequency classes (separated by two vertical dashed lines). U stands for CN-unaltered, G for CN-gain and L for CN-loss. (**b**) Circos diagram showing the chromosomal distribution of different CNV serial orders. The twenty-six different orders are arranged from the rim, showing chromosome banding, inward in accordance to their left-to-right arrangement in Part **a**, *viz.* ULU (blue) in the outermost ring followed by LUU (red), UGU (blue) *etc.*, where orders with a CN-status of U at N-stage are colored blue, and orders with a CN-status of G or L at N-stage colored red. Classes I-III are separated by dashed circles in black. (**c**) GUG/GUL and LUL/LUG frequency ratios determined using different window sizes. Without lineage effects, these ratios would be expected to yield Q=1 (as represented by the red dashed line). In contrast, all the Q-values of GUG/GUL (blue) and LUL/LUG (red) observed in sequence windows varying from 50-kb to 500-kb significantly exceeded unity (*, *p* < 10^-16^). (See Fig. 1 for abbreviations, B, N, P and T)

The plausible basis for these different classes could be straightforward: the CNV orders in Class I entailed minimal copy-number departures from the starting B-stage, and were therefore well tolerated; in comparison, the Class II of CNV orders incurred greater departures from U and were less well tolerated. Every CNV order in the Class III involved at least one double-dose change jumping either from G to L or from L to G between two successive stages of cancer development, a distinct disadvantage that led to their lowest frequencies.

The double-dose disadvantage explained the low frequencies of GLU, LGU, UGL, ULG, LGL, GLG, LGG, GGL, GLL and LLG, but not the low frequencies of LUG and GUL which fell into Class III even though they did not incur any double-dose copy number changes, in contrast to GUG and LUL which belonged to the more abundant Class II. The contrast indicates that lineage effects were important not only to LOH-partitions (Figs. 1b, 3b and 4b), but also to the frequencies of different CNV orders. In GUG, the G status of T-stage cells constituted a reversion to the G status of N-stage cells. In LUL, the L status of T-stage cells likewise constituted a reversion to the L status of the N-stage cells. Thus, both these reversions were favored by lineage effects, allowing GUG and LUL to join Class II even though they each incurred two CNV-status changes. In contrast, lineage effects acted against LUG and GUL, because the CNV-status of the T-stage cells in these cases was not a restoration of the CNV-status of the N-stage cells, thereby explaining their diminished frequencies. These lineage effects were observable when different sizes of sequence windows were employed for CNV identification: as shown in Fig. 7c, both the quotients (Q-value) of GUG/GUL and LUL/LUG greatly exceeded unity (marked by dashed red line), yielding significant lineage effects of *p* < 10^-16^ for all sizes of sequence windows ranging from 50-500kb.

## Discussion

In the present study, genomic sequence analysis employing the AluScan platform revealed that the paratumor region at <2cm from tumor’s edge and the non-tumor tissue region at >2cm from tumor (see Methods) contained numerous GOH and LOH types of SNVs along with CNVs that pointed to these regions as premalignant or early malignant stages in cancer development despite their apparently normal cell morphologies. Residue-by-residue tracing of these mutations through the B-, N-, P- and T-stages pointed to much higher LOH rates and GOH-m rates than GOH-M rates (Fig. 1a) in accord with our earlier findings^17^. In addition, prevalence of FR mutation cycles was observed with strong lineage effects in the partition of LOHs between MM and mm products (Fig. 1b), and in the steeply unequal CNV frequencies between GUG and GUL and between LUL and LUG (Fig. 7c). These findings based on the BN-P-T tetra samples, and supported by those from the B-N-T trio samples (Fig. 3), the N-T-M trio samples based on AluScan, WES and WGS (Figs. 4 and 5) and ICGC B-T paired samples based on WGS (Fig. 6) pointed to a cancer development sequence in the order of B-N-P-T, as represented in the Stage Specific Population (SSP) Model in Fig. 8a, in accord with the greater distance of the N-region than the P-region from the tumor. In this SSP model, while each of the N-, P- and T-stage cell populations could comprise multiple cell clones, a majority of the cell clones within the same stage would display largely similar mutational and morphological characteristics.

**Figure 8.**
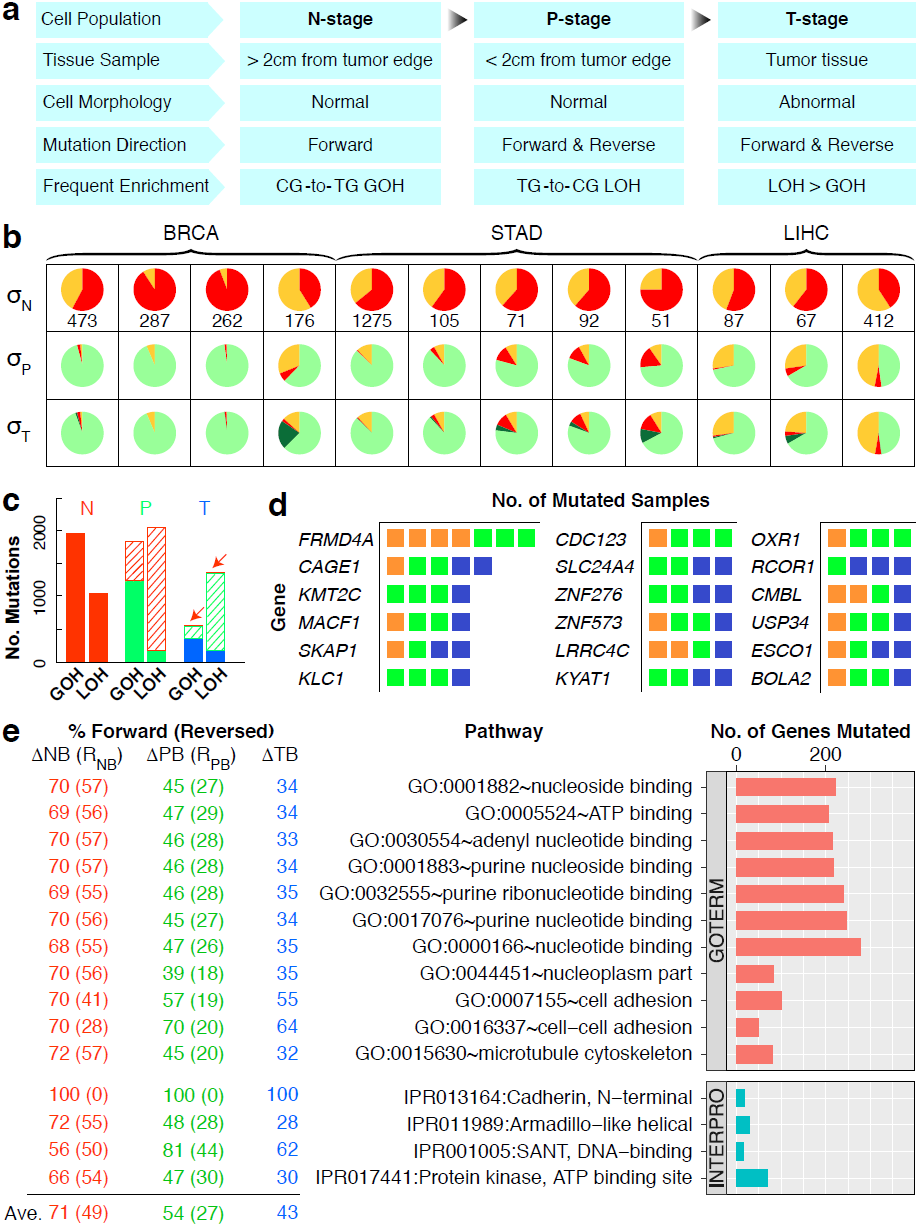
Mutational features of different stages of cancer development. (**a**) Stage Specific Population (SSP) model of cancer development. (**b**) Reversal of N-stage SNVs at P- and T-stages in twelve tetra-sample cases of BRCA, STAD and LIHC. Colored sectors represent N-stage GOHs (red), N-stage LOHs (orange), P-stage reversals of N-stage SNVs (light green), and T-stage reversals of N-stage SNVs (dark green), respectively. σ_N_ shows the total number of SNVs occurred at N-stage compared to B-stage cells with relative proportions of GOH and LOH shown by colored sectors. σ_P_ and σ_T_ show the proportions of N-stage SNVs unreversed and reversed at P-stage and T-stage, as color-coded sectors. For N-stage SNVs, P-stage reversion significantly exceeded T-stage reversion (*p* = 3.1×10^-12^). (**c**) Forward and reverse SNVs in N-, P- and T-stage samples. *Left*: N-stage SNVs comprising forward GOHs and LOHs (solid red). *Middle*: P-stage SNVs comprising forward GOHs and LOHs (solid green); and GOHs and LOHs that reversed N-stage LOHs and GOHs respectively (striped red). *Right*: T-stage SNVs comprising forward GOHs and LOHs (solid blue); and GOHs and LOHs that reversed P-stage LOHs and GOHs (striped green), and N-stage LOHs and GOHs (striped red highlighted by arrows), respectively. See Supplementary Table 3 for data used in this plot. (**d**) Frequently mutated genes in the twelve tetra-sample cases. Each mutation found in a BRCA (orange), STAD (green) or LIHC (blue) sample is represented by a square listed with the gene. (**e**) Pathway enrichments of mutations in the twelve tetra-sample cases. The horizontal bars showing numbers of mutated genes are arranged from top down in the order of increasing Bonferroni corrected *p*-values for pathways in GOTERM and INTERPRO databases, respectively. The % of genes in the pathway that displayed one or more forward SNVs at N-, P- or T-stage is indicated under ΔNB, ΔPB and ΔTB respectively. The % of N-stage SNVs that were reversed subsequently at P- or T-stage was indicated by (R_NB_). The % of P-stage SNVs that were reversed subsequently at T-stage was indicated by (R_PB_). (See Supplementary Tables 12 and 13 for mutated genes and pathways, and Fig. 1 for abbreviations)

Fig. 8b shows that although the twelve B-N-P-T tetra-sample cases were derived from three types of solid tumors, they all displayed a largely similar mutational profile: the SNVs found at N-stage (σ_N_) comprised different proportions of GOHs (red) and LOHs (orange), and underwent substantial reversions in P-stage cells, followed by a much smaller number of additional reversions occurring in the T-stage cells. Altogether, that more N-stage SNVs were reversed in P-stage than in T-stage in all twelve samples amounted to a highly non-random observation (*p* = 3.1×10^-12^), in confirmation of the stage-specific difference in mutational activities between P- and T-stage cells. Overall, the numbers of forward SNVs decreased in P-stage compared to N-stage, and decreased even further in T-stage. The forward N-stage GOHs and LOHs underwent strong reversals in the P-stage, and far less so in T-stage. On the other hand, the forward P-stage GOHs and LOHs were strongly reversed in T-stage (Fig. 8c). The huge difference between the numbers of forward N-stage GOHs and LOHs reversed in P-stage and the numbers of forward N-stage GOHs and LOHs reversed in T-stage clearly attested to the highly dissimilar mutational activities of the N-, P- and T-stage cells. On account of the importance of microenvironment in carcinogenesis^6^, such stage-specific characteristics could arise at least in part from the sharing of a common microenvironment by all cell clones within the same N-, P- or T-stage. The finding of an N-stage cell population characterized by high GOH type of SNVs and normal cell morphology (Fig. 8a) was consisted with the elevated SNV prevalences by 27-fold (*p* < 0.001) or 36-fold (*p* < 0.0001) observed in normal kidney cortices of subjects that were smokers or exposed to the environmental carcinogen aristolochic acid respectively^28^, and pointed to the propensity of N-stage cells to carcinogenesis.

The pronounced reversions of N-stage SNVs in P-stage, P-stage SNVs in T-stage, and T-stage SNVs in M-stage (Figs. 1b and 3b) suggest that FR-cycle between successive development stages could be a common cellular evolution strategy for adjustment to the microenvironmental changes encountered during stage transition. In this regard, it is notable that, when a deviant *Drosophila melanogaster* population induced by extreme starvation was allowed to readapt to the ancestral culture environment, reversions of SNPs back to ancestral allele genotypes over fifty generations of evolution amounted to about 50%^29^, comparable to the average level of SNV-reversions exhibited by P-stage cells in the form of Type-IVa changes in Fig. 2c.

In any event, the wide occurrence of FR-cycles over the N-, P- and T-stages suggests that these stages of cancer development could be burdened with excessive deleterious mutations that require alleviation through reverse mutations to ensure optimal growths. This could help explain the findings of transformed cells multiplying more slowly at low population density than untransformed cells, *e.g.*, upon transformation of a C3H/10T1/2CL8 fibroblast cell line derived from C3H mouse embryos by 3-methylcholanthrene, the transformed cells exhibited a saturation density 2-3 times that of untransformed cells, but generation times of 22 and 27 hours, *viz.* 40-70% longer than the 15.5 hours for the untransformed cells^30^. Likewise, NIH 3T3 cells displayed retarded growth at low density and increased saturation density preceding the formation of transformed loci^31^, while the increased density attained might stem from reduced contact inhibition, and the longer generation times could be the result of excessive deleterious mutations.

The *FRMD4A* gene^32^ was mutated in four BRCA cases and three STAD cases in the twelve tetra-sample cases, and *CAGE1* for cancer antigen 1^33^ was mutated in BRCA, STAD as well as LIHC samples. SNVs recurrent in four out of twelve cases were detected for sixteen different genes (Fig. 8d). Based on the GOTERM and INTERPRO databases, pathway enrichment analysis show that SNVs in the B-N-P-T tetra samples were frequent in the cell-adhesion and protein kinase pathways (Fig. 8e). It was striking that all of the N-stage mutations in the cadherin N-terminal domain family persisted un-reversely throughout the P- and T-stages, pointing to the importance of this family of cell adhesion molecules at multiple stages of cancer development.

In conclusion, because cancers are driven by mutations, the nature of the mutations in the evolving cancer cells furnishes an appropriate basis for delineating the major stages of carcinogenesis. In the present study, the mutational profiles of the cell populations in the N-, P- and T-stage samples showed that N-stage cells harbored large numbers of SNV mutations, more GOHs than LOHs, which were enriched with NCG>NTG type of GOHs with associated CNVs. The P-stage cells displayed, relative to N-stage cells, more LOHs than GOHs. A major fraction of their LOHs represented reversals of the forward GOH mutations found in N-stage, and were enriched with NTG>NCG type of LOHs with associated CNVs. In the T-stage cells, the ratio between LOHs and GOHs was even higher than P-stage cells. At T-stage, there were numerous reversals of P-stage mutations but far fewer reversals of N-stage mutations. Such distinct mutational profiles of the different stages gave rise to FR-cycles of mutational changes between N- and P-stages and between P- and T-stages in a sequential SSP model of cancer development that was supported by pronounced lineage effects on the partition of LOHs between MM and mm products (yellow highlighted in Figs. 1b, 3b and 4b), as well as the relative frequencies of different orders of copy number changes between the N-, P- and T-stages (Fig. 7a), which could not be explained by early divergence of different cell clones that subsequently engaged in parallel evolution. While mutations increased diversity and contributed to the formation of stage specific cell populations, directional selection made evident by MM-over-mm preference in LOHs stemming from germline heterozygous sites and the reversal of SNVs and CNVs to their original genotypes, likely on account of the optimization of cell growth and multiplication, was essential in shaping the distinctive mutational landscapes of different stages of cancer development. In this basis, the possibility arises that early N- and P-stage cells which have not yet accomplished their requisite mutation reversals might be relatively deficient in growth and replication vigor, in which case it could be advantageous to target therapeutic interventions at these early stages of cells before they have accomplished their mutation reversals to become fully malignant, therapy-resistant cancers.

## Methods

### Tumor purity and histology

Tumor purity in all B-N-P-T tetra and B-N-T trio samples was estimated using VarScan software^34^ and ‘absCNseq’ R package^35^. The ‘my.res.list’ function of absCNseq was applied with the following parameters: alpha.min = 0.2, alpha.max = 1, tau.min = 1.5, tau.max = 5, min.sol.freq = 0, min.seg.len = 0, qmax = 7, lamda = 0.5.

For histological and immunohistochemical staining (Fig. 1c and Supplementary Table 1), samples were taken from the tumor, the adjacent paratumor region (< 2cm from tumor), and the non-tumor region (> 2cm from tumor) of a BRCA patient. The samples were fixed in 4% paraformaldehyde, dehydrated, embedded in paraffin, sectioned and subjected to standard hematoxylin and eosin (HE) staining. Immunohistochemical staining for estrogen receptor (ER), progesterone receptor (PR) and human epidermal growth factor receptor-2 (HER2) were conducted following the conventional procedures as described^36^.

### DNA samples for AluScan sequencing

Written informed consent was obtained from each patient who participated in this study. Subject recruitment and sample collection were approved by the institutional ethics review boards of Hong Kong University of Science and Technology, Second Xiangya Hospital of Changsha, Chinese University of Hong Kong, Second Military Medical University of Shanghai, First Hospital of Nanjing, Jiangsu Cancer Hospital, and University of Hong Kong. DNA extraction and AluScan sequencing library preparation were performed as described previously^17,37^. White-blood-cells were treated as representative of germline controls in keeping with the recommendation by the Cancer Genome Atlas (TCGA) project^38^. The N- and P-stage tissues included and subjected to AluScan sequencing in this study were obtained as follows: N-stage tissue was collected at > 2cm from the edge of tumor in the vicinity of tumor, and P-stage tissue was collected at < 2cm from the edge of tumor. The AluScan cancer cases, designated as B-N-P-T, B-N-T or N-T-M sample-sets, were listed in Supplementary Table 1 with demographical and clinical information.

### AluScan sequencing

AluScans of genomic regions flanked by Alu repetitive sequences were obtained by means of inter-Alu PCR as described^17,37^, employing both Head-type and Tail-type Alu consensus-based primers to ensure capture of a vast number of inter-Alu amplicons. In brief, a 25-µl PCR reaction mixture contained 2 µl Bioline 10× NH4 buffer (160 mM ammonium sulfate, 670 mM Tris-HCl, pH 8.8, 0.1% stabilizer; http://www.bioline.com), 3 mM MgCl_2_, 0.15 mM dNTP mix, 1 unit Taq polymerase, 0.1 µg DNA sample, and 0.075 µM each of the four following Alu consensus sequence-based PCR primers:

AluY278T18 (5’-GAGCGAGACTCCGTCTCA-3’);

AluY66H21 (5’-TGGTCTCGATCTCCTGACCTC-3’);

R12A/267 (5’-AGCGAGACTCCG-3’);

L12A/8 (5’-TGAGCCACCGCG-3’).

PCR was carried out at 95 °C, 5 min for DNA denaturation, followed by 30 cycles each of 30 s at 95 °C, 30 s at 50 °C, and 5 min at 72 °C, plus finally another 7 min at 72 °C. Amplicons were purified with ethanol precipitation, sequenced on the Illumina HiSeq platform at Beijing Genomics Institute (Shenzhen, China) and mapped to the hg19 reference human genome for downstream bioinformatic analysis.

### WGS and WES raw data

Whole-genome sequencing (WGS) data generated from tumor-blood paired samples with the Illumina system by the International Cancer Genome Consortium (ICGC) and the Cancer Genome Atlas (TCGA) were downloaded in fastq format with permission (https://www.synapse.org/#!Synapse:syn2887117). These included the Pilot-63 set, and eighty-six hepatocellular carcinoma (LIHC), seventy-five non-small-cell lung cancer (NSCLC) and twenty-two intrahepatic cholangiocarcinoma (ICC) cases (Supplementary Table 10) with information accessible through the ICGC Data Portal (https://dcc.icgc.org). In addition, raw WGS data generated by Ouyang *et al*^20^ from four hepatitis B positive LIHC patients having pulmonary metastasis were obtained along with data from same-patient liver tissue controls, and included in the N-T-M trio sample analysis as the WGS-Liver-M subset. Moreover, raw data of whole-exome sequencing (WES) from sixty-seven brain metastatic cancer patients were obtained from Brastianos *et al*^21^. Tissues sampled at > 2cm from the edge of tumors were used as normal control tissues in Ouyang *et al*^20^ and Brastianos *et al*^21^, and treated as non-tumor, or N-stage, samples in this study.

### SNV calling

For the paired-end sequencing reads generated on the Illumina platform by the AluScan, WGS or WES methods, bioinformatics analysis including alignment, sorting, recalibration, realignment and removal of duplicates using BWA (Burrows-Wheeler Aligner, version 0.6.1)^39^, SAMtools (Sequence Alignment/Map, version 0.1.18)^40^ and GATK (Genome Analysis Tool-Kit, version 3.5) were performed for identification of single nucleotide variations (SNVs) according to the standard framework^41^ as described previously^17,37^. The ‘UnifiedGenotyper’ module of GATK was employed for genotyping of SNVs. Only genomic sequence regions with enough coverage, *i.e.*, read depth > 8, were included in the analysis, and the following parameters were applied to filtrate for SNVs of different genotypes: major allele frequency ≥ 95% for the ‘MM’ loci; major allele frequency ≥ 30% and ≤ 70%, and QD ≥ 4 for ‘Mm’ or ‘mn’ loci; and minor allele frequency ≥ 95% and QD ≥ 20 for ‘mm’ loci. Strand bias estimated using Fisher’s Exact Test (FS) was employed to ensure FS value ≤ 20 for both heterozygous ‘Mm’ or ‘mn’ loci and homozygous ‘MM’ or ‘mm’ loci.

For cancer cases with more than two samples from each patient, *i.e.*, the B-N-P-T tetra-sample set of twelve cases, the B-N-T trio-sample set of seventeen cases and the N-T-M trio-sample set of seventy-three cases, the above mentioned calling of SNVs were first performed for each sample of each case in the multiple sample sets. For each of the multiple-sample cases, only nucleotide positions conformed to all the above SNV calling criteria in every samples of the same patient were included in further analysis. Sites not covered in further analysis were arising from either lack of sequencing reads or failure to meet filtering criteria in any one of the samples of the same patient.

### Mutational profiles of SNVs

Mutational profiles of SNVs were analyzed following the procedure developed by Alexandrov *et al*^18^. For each SNV site, the SomaticSignatures package^42^ under R environment was employed to determine its preceding and following bases. The results were unnormalized for the observed trinucleotide frequencies in the human genome. The resulting mutation frequency profiles were illustrated in three different graphical presentations, *i.e.*, the alteration-group plot, context-group plot and mutation-rate diagram. Custom R scripts for drawing the three different presentations are available at GitHub website (https://github.com/hutaobo/ProfilePlots).

### CNV calling and identification of recurrent CNVs

From AluScan data, the AluScanCNV software^43^ was employed to call paired CNVs between B- and N-stage (ΔNB), between N- and P-stage (ΔPN), between P- and T-stage (ΔTP), between B- and P-stage (ΔPB), as well as between B- and T-stage (ΔTB) samples of the same patient in the B-N-P-T tetra-sample sets of twelve cases, using fixed window sizes ranging from 50- to 500-kb. The ΔNB, ΔPB and ΔTB were arranged sequentially to yield the twenty-six possible serial orders shown in Figs. 7a and 7b. To identify recurrent CNVs, all CNVs found in any sequence window of any of the twelve tetra-sample cases at any stages, including ΔNB, ΔPN and ΔTP, were aggregated. CNVs located in the recently identified Distal Zones^22^ were removed from further analysis to reduce background noise introduced by less informative windows in the human genome. Only sequence windows where CNV was detected in six or more of the twelve patients were considered to harbor a recurrent CNV.

### Co-localization of CNVT and CpGe/MeMRE

CpGe and MeMRE entries were downloaded from UCSC Genome Browser as described^22^, and somatic CNV (CNVT) entries classified as “Copy Number Variants” were downloaded from COSMIC database (http://grch37-cancer.sanger.ac.uk/cosmic/download). The human genome was divided into tandem 2000-bp windows, and the average densities of CNVT breakpoints and base pairs in CpGe or MeMRE in each window were calculated. Thereupon the windows with zero CpGe or MeMRE density were removed to avoid error caused by missing data, and the remaining windows were separated into ten groups based on the percentile of CpGe or MeMRE density. Finally, the average CNVT breakpoint densities in the groups were plotted against the percentile CpGe or MeMRE density.

### Mutation enrichment in genes and pathways

The results of variant analysis of AluScan data of the twelve tetra-sample cases were uploaded to BioMart of the Ensembl database to generate a list of their gene contents under R environment using the ‘biomaRt’ R package^44^. For the ‘getBM’ function, ‘chromosome_name’,‘start_position’,‘end_position’,‘external_gene_name’,‘ensembl_gene_id’and‘description’ were selected as attributes, with ‘chromosomal_region’ filter type, ‘sublist’ filter value, ‘ENSEMBL_MART_ENSEMBL’ biomart type, and ‘grch37.ensembl.org’ host. The resultant gene list was uploaded to DAVID Bioinformatics Resources 6.7^45^ using ‘Functional Annotation Tool’ to obtain three lists of mutation enriched functional groups and pathways as annotated in the three databases GOTERM, InterPro and KEGG, respectively, with mutated genes specified for each groups and pathways. Only those functionally annotated groups and pathways yielding Bonferroni corrected *p*-value, Benjamini corrected *p*-value as well as FDR *q*-value all less than 0.05 were considered statistically significant.

### Statistical analysis and data visualization

Statistical analyses were performed using R software (http://www.r-project.org). The significance probability (*p*) values were calculated by the two-tailed t.test or chisq.test functions in R, and the Pearson correlation coefficients (*r*) were calculated by the cor function in R. Figures were drawn using the ggplot2, lattice or ellipse package under R environment, except for Fig. 7b which was drawn using the Circos program^46^.

## Disclosure of potential conflicts of interest

The authors declare no competing financial or non-financial interests.

## Author contributions

HX conceived and initiated the study; SJD, YL, LC, JFC, RY, AK, GKKL, DHZ, EXZ, LX, WSP and HYW organized and collected the clinical samples and data; TH, YK, IS, WKM, ZW, XL, CHC, PL, SKN, TYCH, JY, XD, SYT and XZ analyzed the samples and data; and HX, TH and SYT wrote the paper. The International Cancer Genome Consortium (ICGC) which led a worldwide cancer genomics collaboration including a project on pan-cancer analysis of whole genomes (PCAWG), contributed tumor-control sample pairs collected and sequenced by PCAWG to the present study.

## Acknowledgments

The study was supported by grants to H. Xue from University Grants Council of Hong Kong SAR (ITS/113/15FP, VPRDO09/10.SC08, VPRDO14SC01, DG14SC02, SRFI11SC06 and SRFI11SC06PG), and grants to J. F. Chen (National 973 Basic Research Program of China, No. 2013CB911300; National Natural Science Foundation of China, No. 81272469; and Natural Science Foundation of Jiangsu Province special clinical project, No. BL2012016). Y. Kumar was a recipient of International Ph.D. Studentship from Hong Kong University of Science and Technology. X. Long was a recipient of Hong Kong Ph.D. Fellowship from Government of Hong Kong SAR. F.W. Pun was recipient of Research Fellowship from HKUST Jockey Club Institute of Advanced Study. Hututa Technologies Limited assisted with the computation facilities.

## Supplementary Information

**Supplementary Table 1**. Information on 105 samples analyzed by AluScan.

**Supplementary Table 2**. Tumor purities of B-N-P-T tetra and B-N-T trio samples estimated by absCN-seq.

**Supplementary Table 3**. Summary of SNV mutations in B-N-P-T tetra samples.

**Supplementary Table 4**. The exact residue-by-residue SNV mutations in each sample of the B-N-P-T tetra-sample cases.

**Supplementary Table 5**. Summary of CNV mutations in B-N-P-T tetra samples.

**Supplementary Table 6**. Summary of SNV mutations in B-N-T trio samples.

**Supplementary Table 7**. The exact residue-by-residue SNV mutations in each sample of the B-N-T trio-sample cases.

**Supplementary Table 8**. Summary of SNV mutations in N-T-M trio samples. (**a**) AluScan; (**b**) WES-Non-Lung; (**c**) WES-NSCLC-L; (**d**) WES-NSCLC-H; and (**e**) WGS-Liver-M.

**Supplementary Table 9**. The exact residue-by-residue SNV mutations in each sample of the AluScan N-T-M trio-sample cases.

**Supplementary Table 10**. Numbers of GOHs and LOHs in 22 ICC, 86 LIHC and 75 NSCLC samples.

**Supplementary Table 11**. The exact window-by-window CNV mutations in each sample of the B-N-P-T tetra-sample cases.

**Supplementary Table 12**. List of genes harboring SNV mutations in B-N-P-T tetra-sample cases.

**Supplementary Table 13**. SNV mutations enriched pathways and genes in B-N-P-T tetra-sample cases.

**Supplementary Figure 1**. Total numbers of different dinucleotide sites in the human genome. Numbers of CG as well as other fifteen types of dinucleotides in human reference genome hg19 are plotted out.

**Supplementary Figure 2**. Mutational profiles of Pilot-63 samples from ICGC analyzed by WGS. (**a**) Context-group plots of GOH and LOH types of SNVs. Each plot is arranged by the ten context groups: A.A, C.C, A.G, C.A, A.C, G.A, C.G, A.T, T.A and G.C, designating the different immediate 5’ and 3’ flanking nucleotides. The opposing GOH mutations (left panel) or opposing LOH mutations (right panel) are placed side-by-side, *e.g.*, by pairing the ATG>ACG GOH (pink bar) with the ACG>ATG GOH (blue bar) in Section-3 of left panel, and similarly pairing the ATG>ACG LOH (pink bar) with the ACG>ATG LOH (blue bar) in Section-3 of right panel (marked by arrowheads and color-coded as in Fig. 6c). (**b**) Mutation-rate diagrams for GOHs. Each of the ten diagrams of triplet duplexes correspond to a context group, labeled 1-10 as in Part **a**. The mutation rates of opposing GOH mutations are labeled on double-headed arrows, except for the single-headed curved arrows in groups 7-10, where the two sequences are identical in a triple duplex. Each double-headed arrows is accompanied by two color-coded mutation rates that correspond to the heights of color-coded bars in Part **a**, *e.g.*, in context group 1, the conversion of double-stranded ACA/TGT to AAA/TTT is associated with a mutation rate of 162, colored red to correspond to the red C>A bar with A.A context in the left panel of Part **a**; whereas the opposing conversion of AAA/TTT to ACA/TGT is associated with a mutation rate of 641, colored orange to correspond to the orange A.C bar with A.A context in the left panel of Part **a**. (**c**) Mutation-rate diagrams for LOHs. The arrows employed are similar to those in Part **b**. All arrows in Parts **b** and **c** are shown as dashed lines for transitions (TSs) or solid lines for transversions (TVs). In the ten diagrams in Part **b** or Part **c**, the boxed TS/TV ratio given for each diagram represents the ratio pertaining to all the TS and TV mutations in the diagram, *e.g.*, in Diagram-1 of Part **b**, TS equals the sum of the four TS rates in the diagram, and TV the sum of the eight TV rates, yielding TS/TV = 1430/2804 = 0.51. The different rates in the diagrams in Parts **b** and **c** are color-coded as in Part **a**.

